## Supplementary Information for "Histone dynamics mediate DNA unwrapping and sliding in nucleosomes: insights from multi-microsecond molecular dynamics simulations"

### Detailed simulated systems' description

**NCP**$_{147}$ - based on an X-ray structure of NCP with PDB ID 1KX5.[1] It has 147 bp DNA derived from a human alpha-satellite DNA sequence repeat. Octamer is based on canonical histones of *X. laevis.* DNA sequence forms two symmetric palindromic 73 bp segments in the structure plus one base pair is strictly at the dyad. The system has full-length histone tails. Symmetry related histone tails were put in identical starting conformations.
**NCP**$_{147}^{tt}$ - analogous to NCP$_{147}$, but histone tails are truncated in this system.
**NCP**$_{146}^{tt}$ - based on an X-ray structure of NCP with PDB ID 1AOI.[2] It has 146 bp of palindromic DNA derived from human X-chromosome alpha-satellite DNA repeat. Octamer is based on canonical histones of *X. laevis.* Histone tails are truncated in this system.
**NCP**$_{145}^{tt}$ - based on an X-ray structure of NCP with PDB ID 3LZ0.[3] It has 145 bp of Widom 601 strong positioning DNA sequence. Octamer is based on canonical histones of *X. laevis.* Histone tails are truncated in this system.
**NCP**$_{147}^{fixed}$ - same as NCP$_{147}$, but histone folds' C-$\alpha$-atoms are fixed (restraints of 1000 kJ/mol/nm$^2$).

System parameters are further presented in Table ST1.

### Supplementary Movies

- **Supplementary Movie 1**: Dynamics of NCP$_{147}$ system.
- **Supplementary Movie 2**: Dynamics of NCP$_{147}^{tt}$ system.
- **Supplementary Movie 3**: Dynamics of NCP$_{145}^{tt}$ system.
- **Supplementary Movie 4**: Dynamics of NCP$_{146}^{tt}$ system.
- **Supplementary Movie 5**: Dynamics of NCP$_{147}^{fixed}$ system.

- **Supplementary Movie 6**: Dynamics of contacts between anchor residues and nucleosomal DNA in $NCP_{147}$ system.
- **Supplementary Movie 7**: Dynamics of contacts between anchor residues and nucleosomal DNA in $NCP^{tt}_{147}$ system.
- **Supplementary Movie 8**: Dynamics of contacts between anchor residues and nucleosomal DNA in $NCP^{tt}_{145}$ system.
- **Supplementary Movie 9**: Dynamics of contacts between anchor residues and nucleosomal DNA in $NCP^{tt}_{146}$ system.
- **Supplementary Movie 10**: Dynamics of contacts between anchor residues and nucleosomal DNA in $NCP^{fixed}_{147}$ system.

### Interactive data

Available at `http://intbio.github.io/Armeev_et_al_2021`.

### Supplementary Tables

See table(s) ST1, ST2 below.

Table ST1: Parameters of the simulated systems.

| Name | PDB ID[a] | Box volume, $nm^3$ | Atom number | Water molecules | Ions number, Na/Cl | Ionic strength by water volume, mM | Ionic strength by box volume, mM | Bulk ionic strength[b], mM | Minimum periodic image distance, nm | Time, $\mu s$ |
|---|---|---|---|---|---|---|---|---|---|---|
| $NCP_{147}$ | 1KX5 | 2658 | 267K | 81K | 363/219 | 150 | 136 | 172 | 2.089 | 8.9 |
| $NCP^{tt}_{147}$ | 1KX5 | 2914 | 293K | 90K | 468/242 | 149 | 137 | 168 | 2.53 | 7.4 |
| $NCP^{tt}_{145}$ | 3LZ0 | 2817 | 283K | 87K | 459/237 | 151 | 139 | 168 | 2.47 | 8.6 |
| $NCP^{tt}_{146}$ | 1AOI | 2641 | 265K | 81K | 445/221 | 151 | 138 | 178 | 3.52 | 3.9 |
| $NCP^{fixed}_{147}$ | 1KX5 | 2425 | 245K | 73K | 343/199 | 151 | 136 | 170 | 1.48 | 0.5 |

[a] PDB database ID used to derive the simulated system model.

[b] Calculated at a distance beyond 1 nm from the nucleosome.

[c] See Figure 1e for the location of truncation sites.

Table ST2: Trajectory averaged number of contacts between DNA and histone globular core or histone tails for $NCP_{147}$ system

| Classification method | Contact types | Histone core[a] | Histone tails |
|---|---|---|---|
| All contacts | | 601 | 1219 |
| Nucleotide parts | Phosphate | 458 | 583 |
| | Base | 10 | 288 |
| | Sugar | 133 | 347 |
| Amino acids parts | Backbone | 200 | 466 |
| | Side chain | 401 | 753 |
| Interaction types | Hydrophobic | 22 | 105 |
| | Salt bridges | 34 | 29 |
| | Polar | 66 | 152 |
| | hydrogen bonds | 34 | 263 |
| DNA groove | Major | 1 | 123 |
| | Minor | 9 | 162 |

[a] Core residues are non-tail residues. See tail regions definition in Figure 1e.

### Supplementary Figures

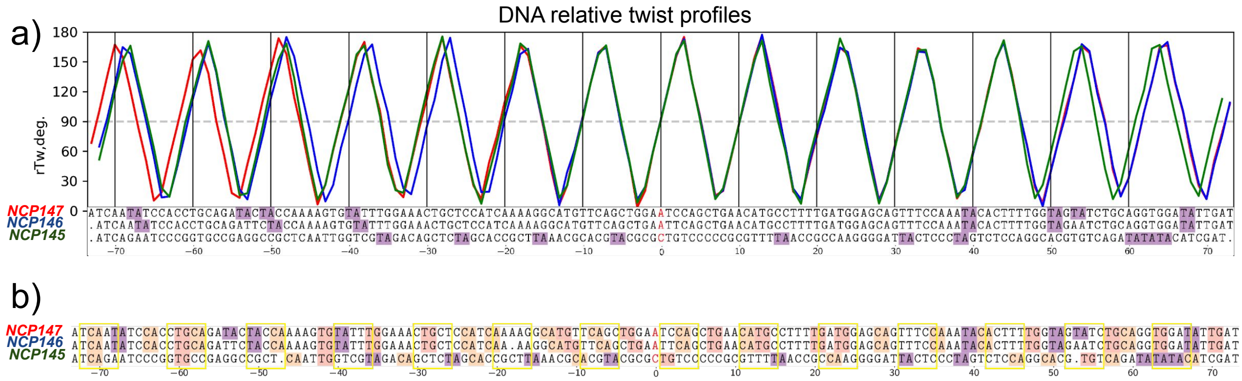


Figure SF1: a) Sequence alignment (by the dyad position) of the DNA sequences (top chain) used in the simulated systems and their relative twist profiles (rTw) in X-ray structures (indicating the orientation of DNA base pairs relative to the octamer surface - see Methods). Red nucleotides mark the dyad position, flexible TA dinucleotides are highlighted in purple; b) Structure based sequence alignment for the top DNA strand (chain I) generated by USCF Chimera.[4] Indels around SHL ±5 are detected (the results for the bottom strand are nearly equivalent). Yellow frames mark regions, where a minor groove faces towards the histone octamer; pyrimidine/purine dinucleotides are highlighted (TA - purple, CA - apricot, TG - pink).

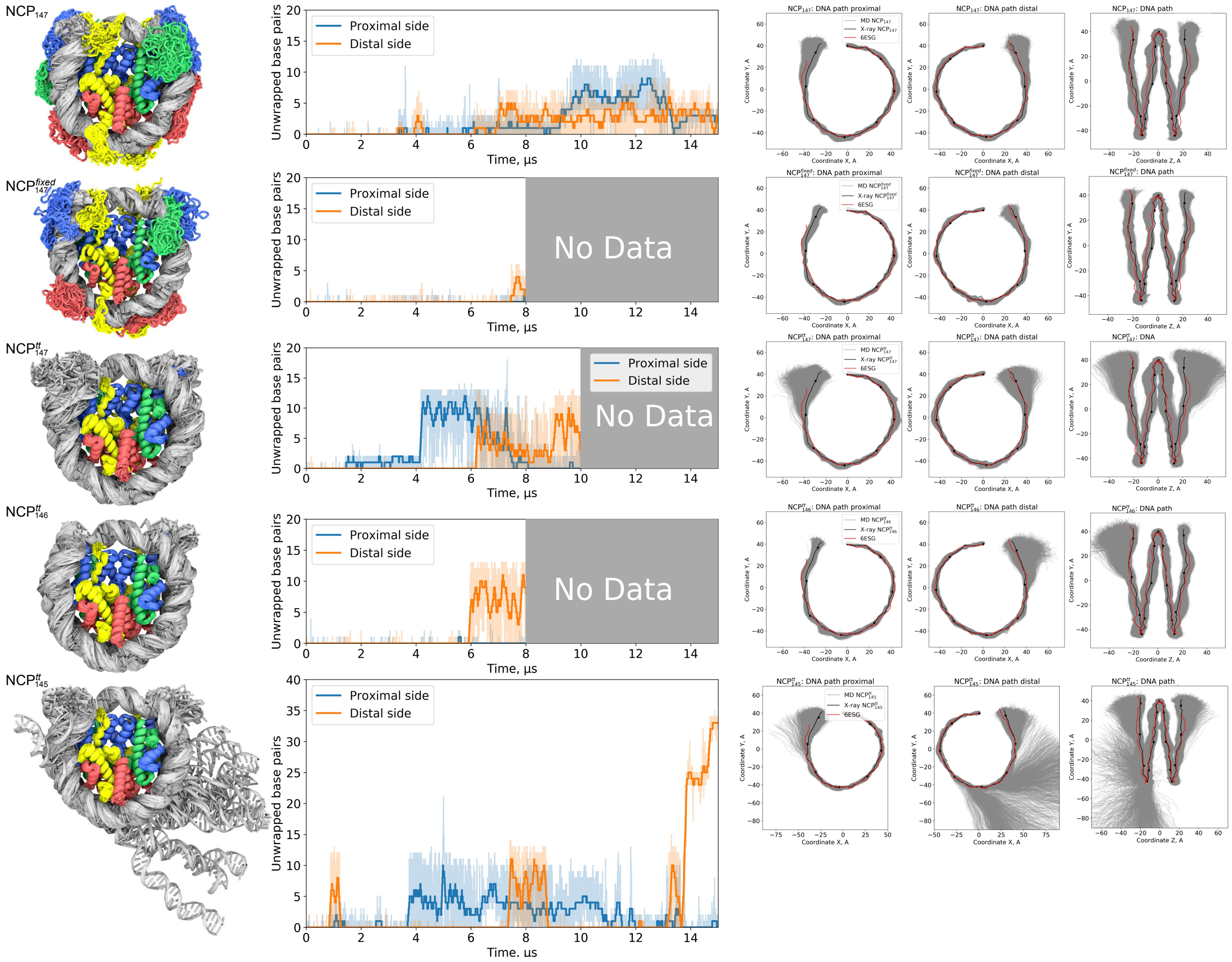


Figure SF2: Overview of the dynamics of the simulated systems. First column: overlay of snapshots along the whole MD trajectory for each system (every 100 ns). Second column: plots of the extent of DNA unwrapping (as estimated by displacement of base pair centers). Right three columns: 2D projections of DNA path (base pair centers) in nucleosome reference frame for frames along the MD trajectory.

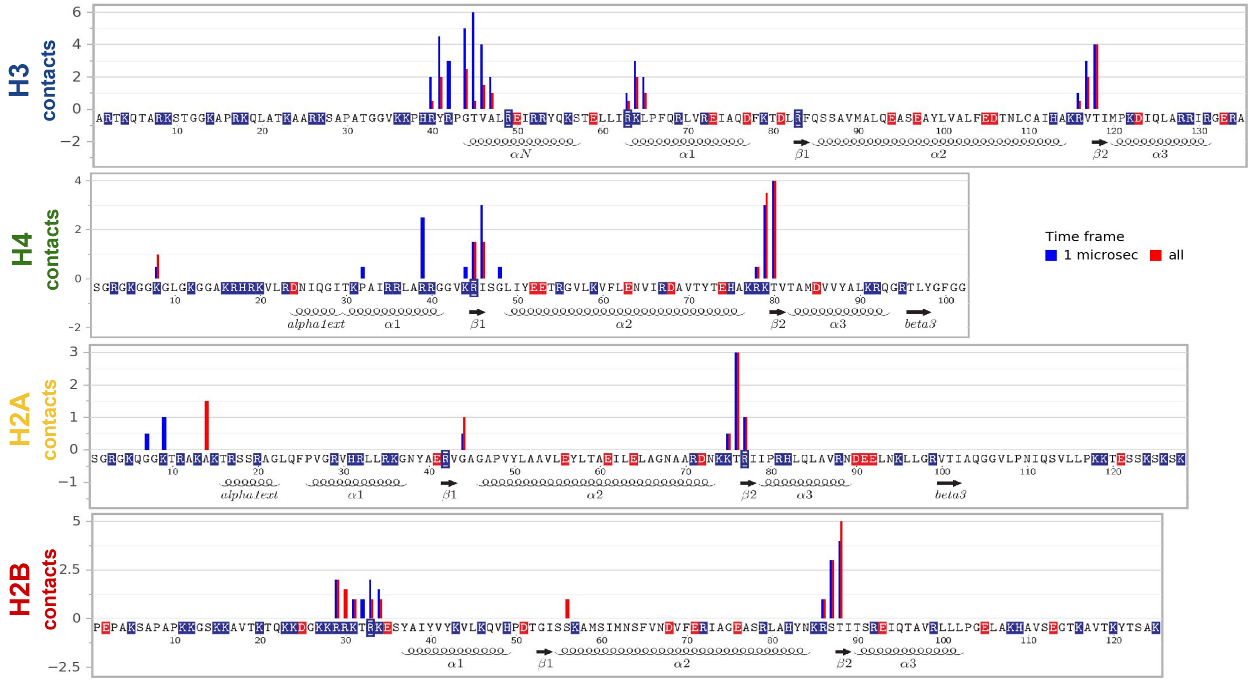


Figure SF3: Number of stable atom-atom contacts between histone residues and nucleosomal DNA plotted along the histone sequences for $NCP_{147}$ simulation. Stable atom-atom contacts are defined as those present in at least 40% of MD frames (the threshold is lower than for stable residue-nucleotide contacts because atom-atom contacts are less persistent). Contacts are averaged over symmetry-related histone chains. The data is shown for the first 1 $\mu$s and the full 15 $\mu$s trajectory.

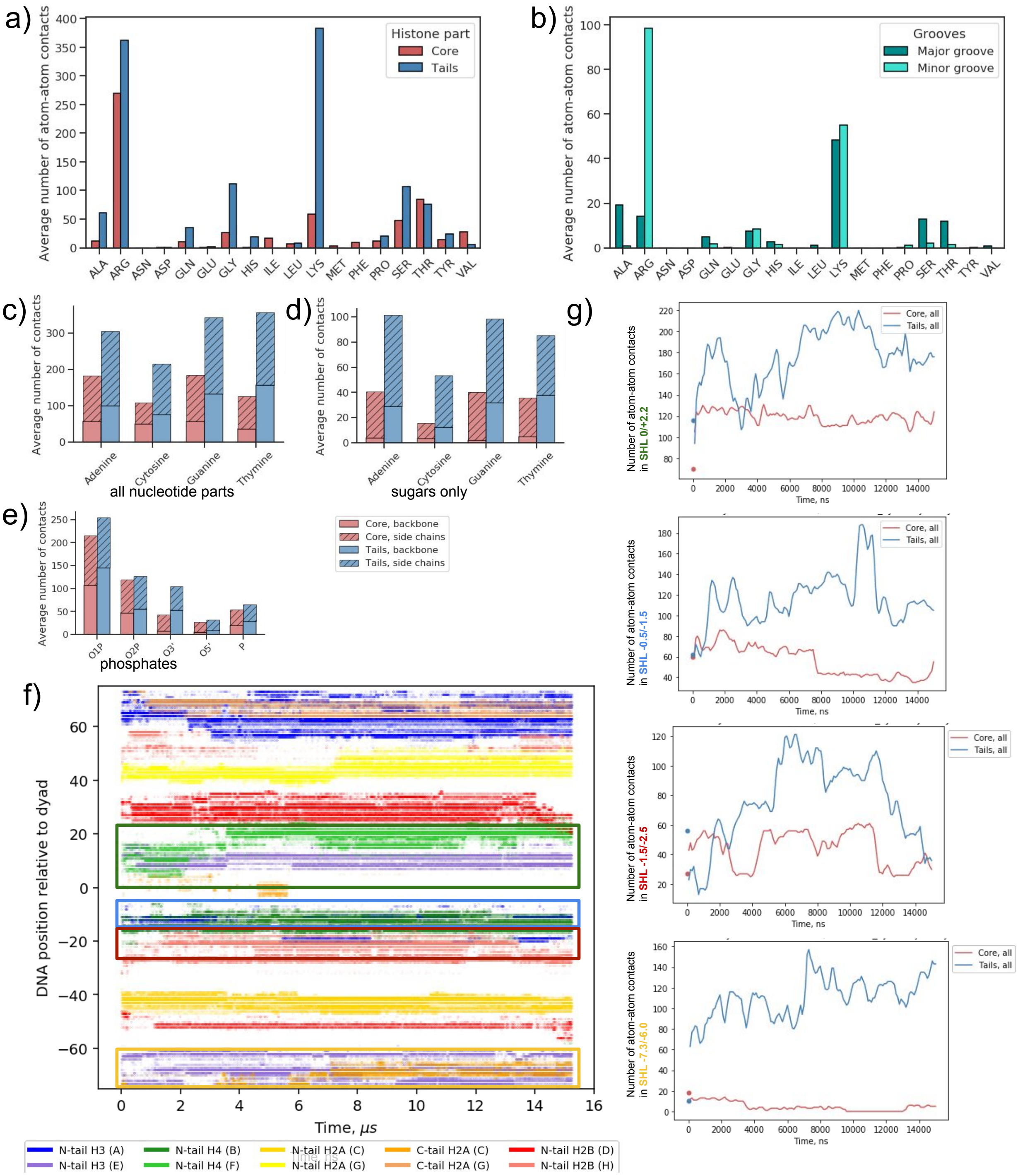


Figure SF4: Evolution and statistics of histone-DNA atom-atom contacts for $NCP_{147}$ simulations. a) Number of contacts classified by histone part and amino acid residue; b) contacts between histones and DNA bases classified by amino acid residue and DNA major/minor groove; c) contacts between histones and DNA classified by nucleotide; d) contacts between histones and DNA sugar moieties classified by nucleotide; e) contacts between histones and DNA phosphates classified by phosphate atoms; f) a map representing the evolution of interactions between histone tails and different positions in nucleosomal DNA; g) evolution of atom-atom contacts between core and tail parts of histones for different segments of nucleosomal DNA defined by their SHLs and highlighted by color boxes in panel f.

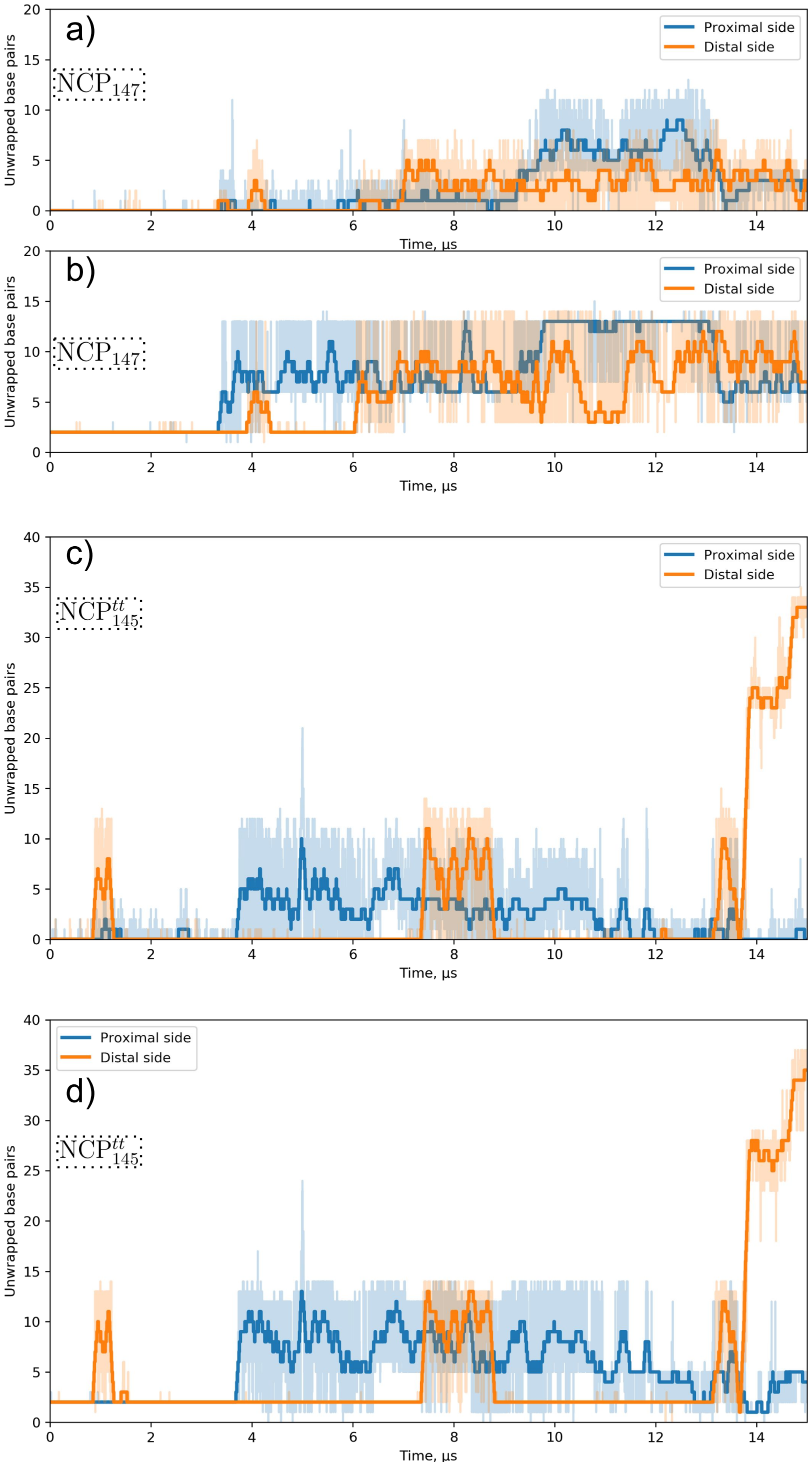


Figure SF5: Comparison of two approaches to quantify DNA unwrapping for $NCP_{147}$ and $NCP^{tt}_{145}$ systems. a),c) distance criterion: displacement of all the base pairs of the segment by more than $7\mathring{A}$ from the DNA path in the corresponding X-ray structure. b),d) contacts criterion: loss of contacts with the globular core (non-tail part) of the histone octamer.

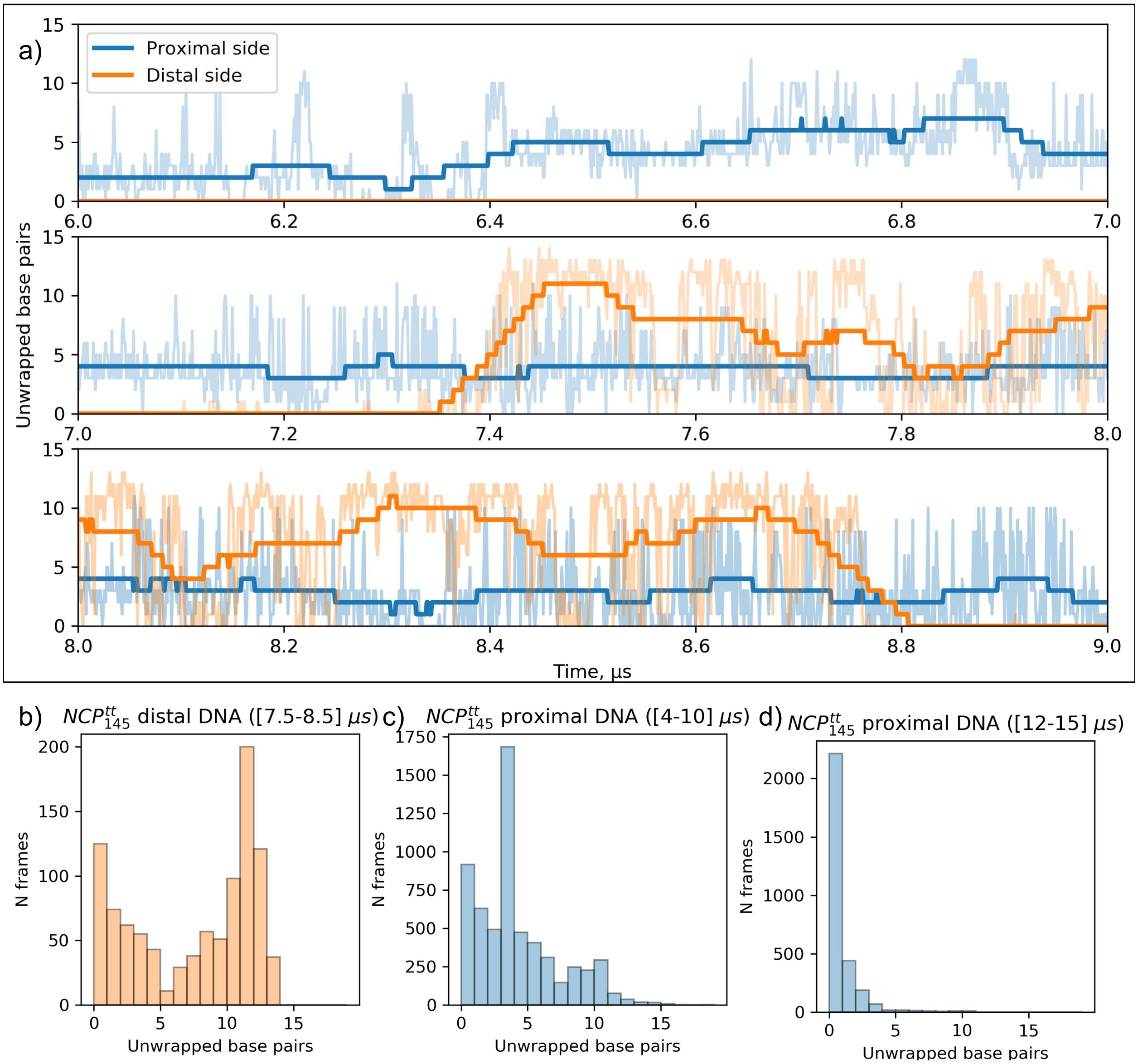


Figure SF6: A detailed view of DNA unwrapping dynamics for $\mathrm{NCP}^{tt}_{145}$ simulation. a) A zoom-up view of DNA unwrapping dynamics between 6-9 $\mu$s. b) Histogram of different unwrap values for the distal DNA end sampled during period 7.5-8.5 $\mu$s. c) Histogram of different unwrap values for the proximal DNA end sampled during period 4-10 $\mu$s. d) Histogram of different unwrap values for the proximal DNA end sampled during period 12-15 $\mu$s.

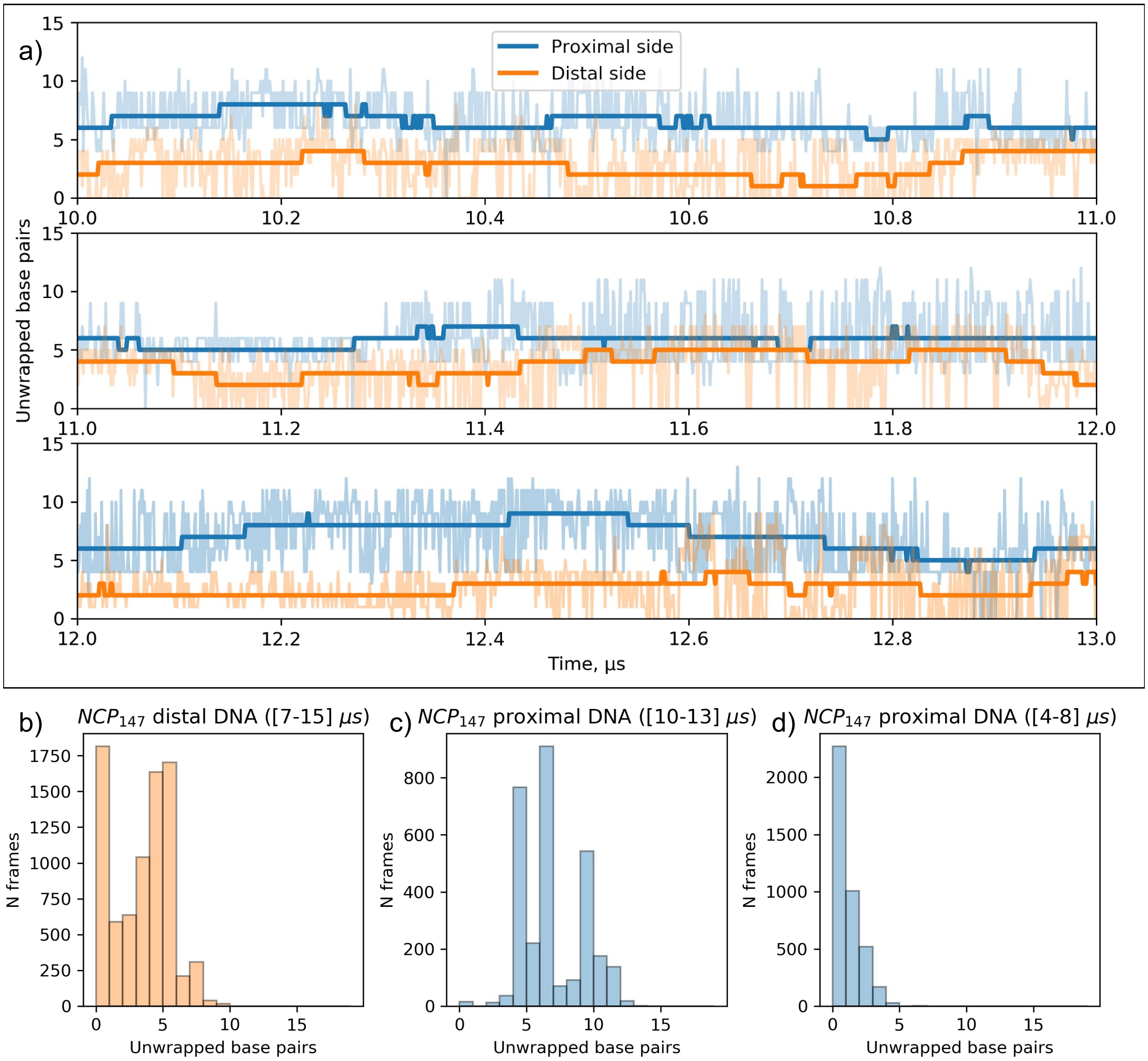


Figure SF7: A detailed view of DNA unwrapping dynamics for $NCP_{147}$ simulation. a) A zoom-up view of DNA unwrapping dynamics between 10-13 $\mu$s. b) Histogram of different unwrap values for the distal DNA end sampled during period 7-15 $\mu$s. c) Histogram of different unwrap values for the proximal DNA end sampled during period 10-13 $\mu$s. d) Histogram of different unwrap values for the proximal DNA end sampled during period 4-8 $\mu$s.

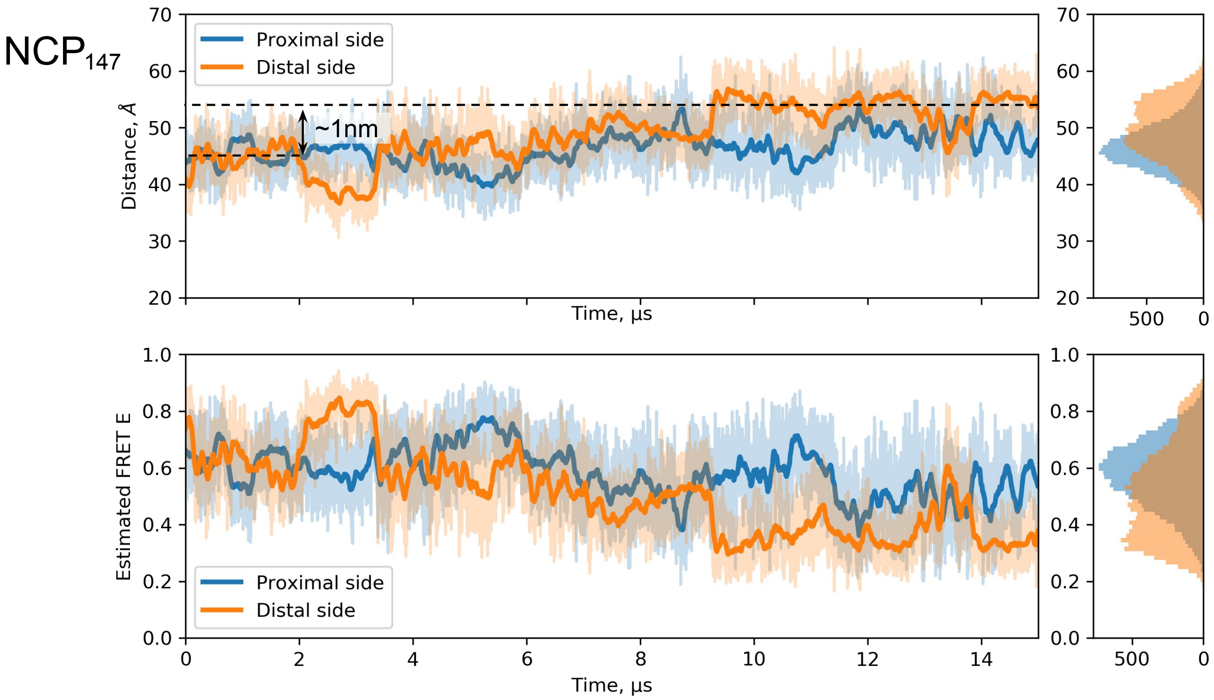


Figure SF8: Distances (top panel) and simulated FRET efficiencies (bottom panel) between FRET labels placed on DNA positions -73 and 2 for the proximal side and -2 and 73 for the distal side following ref.[5] for $NCP_{147}$. We estimated the distances between the fluorescent dyes' attachment sites as the distances between O5′ atoms of the sugar-phosphate backbone of the respective nucleotides. FRET efficiencies were calculated as $E = \dfrac{1}{1+(r/R_0)^6}$ were $r$ is distance between dyes and Förster radius $R_0 = 49\mathring{A}$ as in ref.[5]

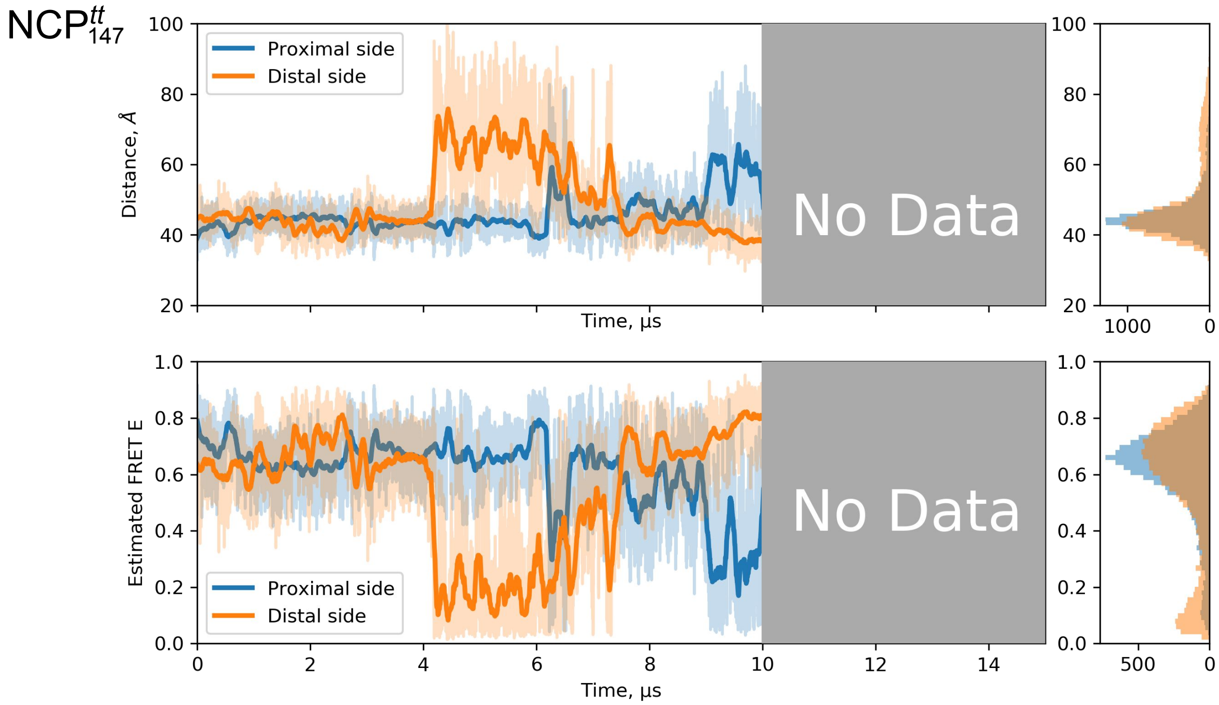


Figure SF9: Distances (top panel) and simulated FRET efficiencies (bottom panel) between FRET labels placed on DNA positions -73 and 2 for proximal side and -2 and 73 for distal side following ref.[5] for NCP$^{tt}_{147}$. Distances and FRET efficiencies are measured as in SF8.

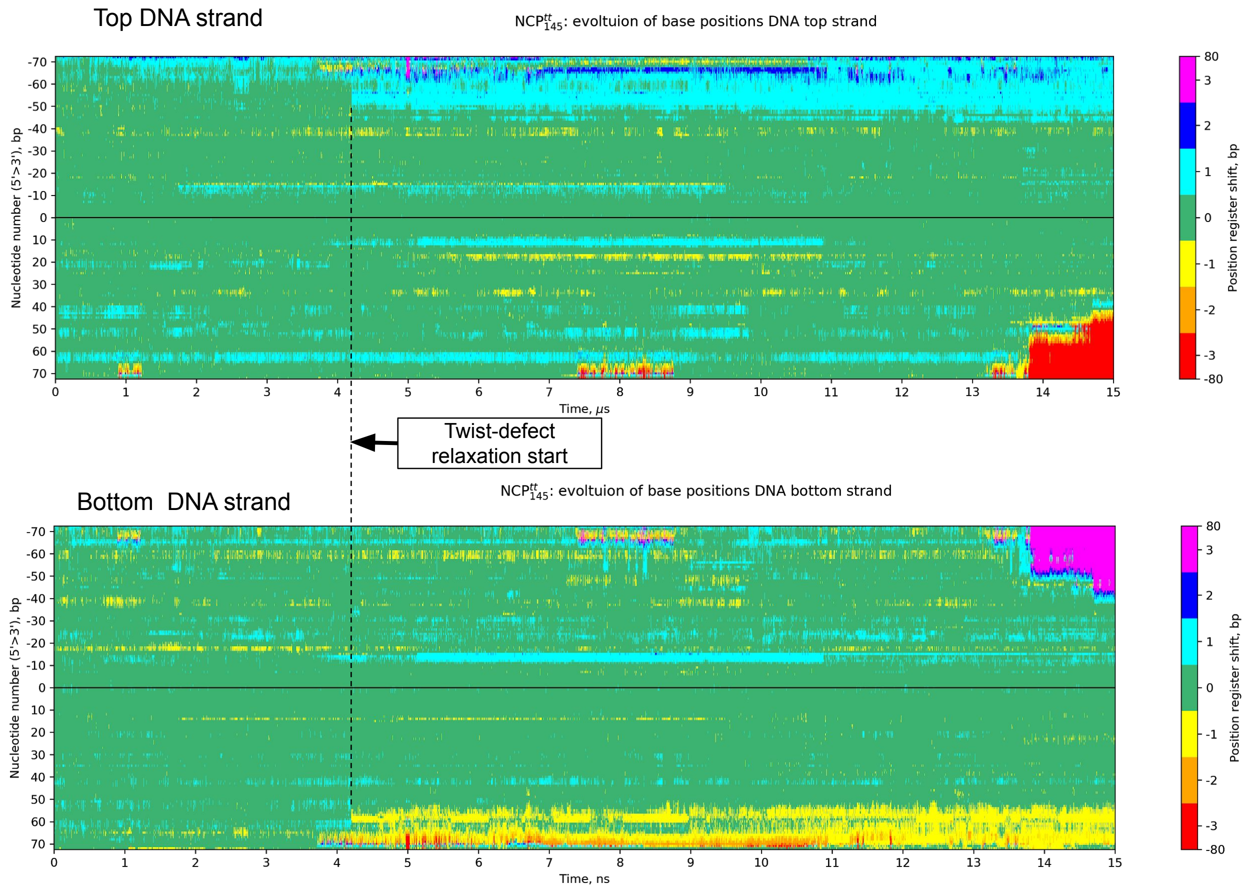


Figure SF10: DNA strand position register shift during simulations of NCP$^{tt}_{145}$. The plot is similar to Figure 7e, but shows data for the full range of the bottom and top DNA strands.

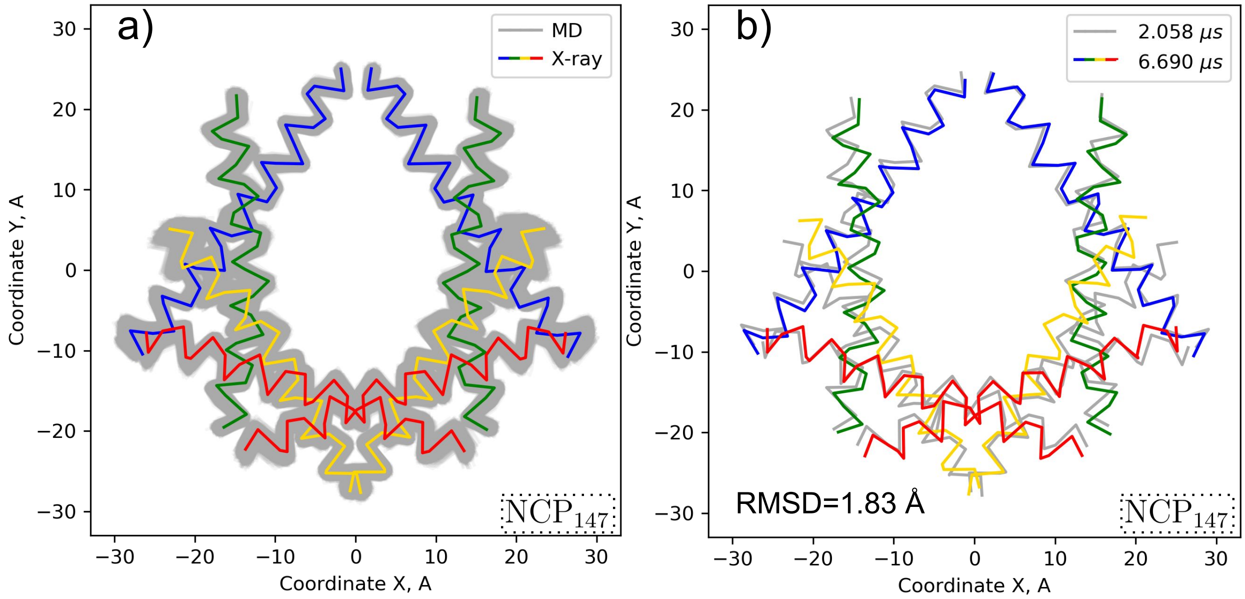


Figure SF11: Analysis of histone octamer plasticity in $NCP_{147}$ simulation. The description is similar to Figure 8

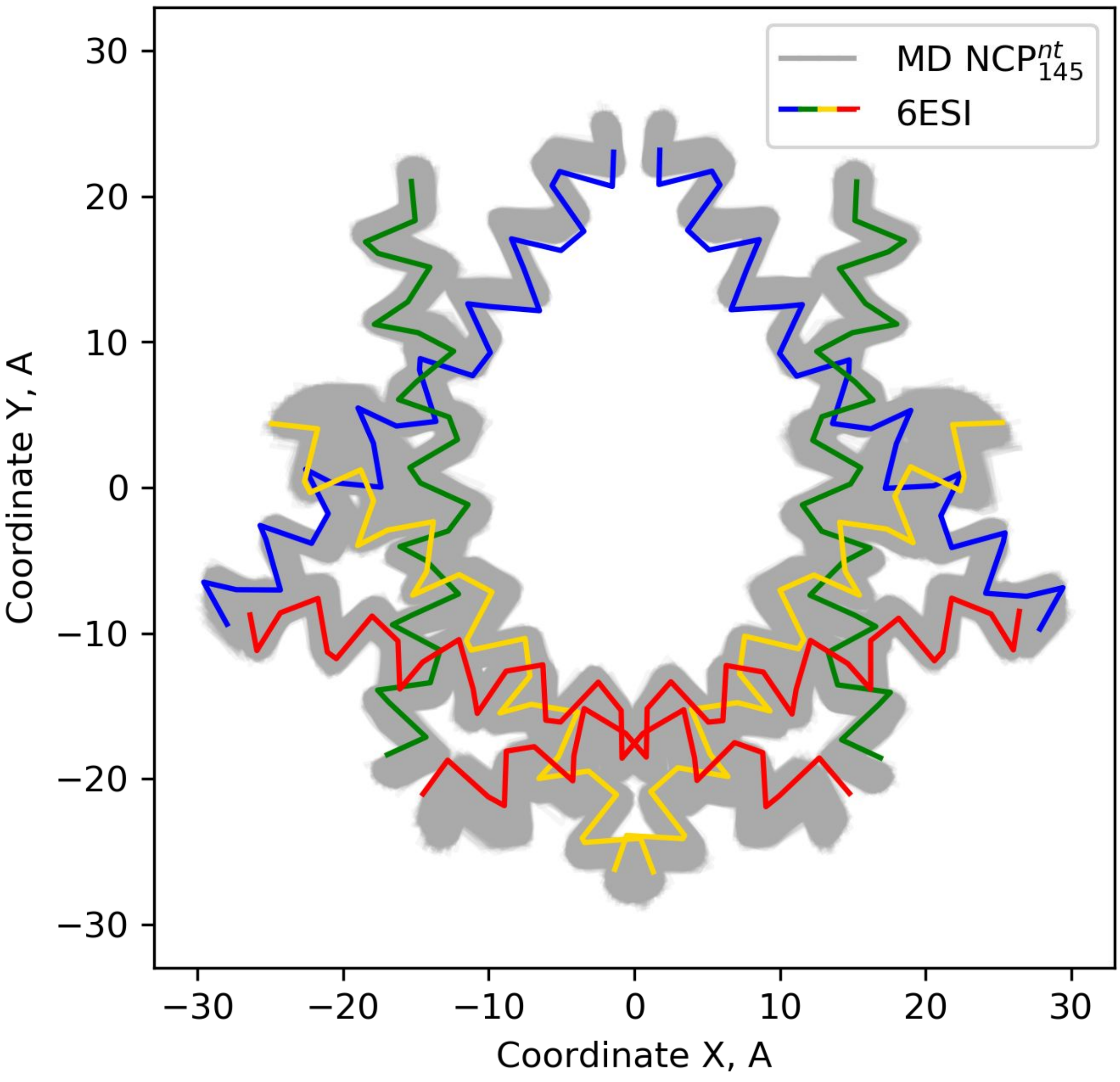


Figure SF12: 2D projections of the histone $\alpha$2-helices from $NCP^{tt}_{145}$ simulation vs. their projections in the recently reported deformed (squeezed by 8% along the dyad) NCP structure seen in cryo-EM (PDB ID 6FQ6).

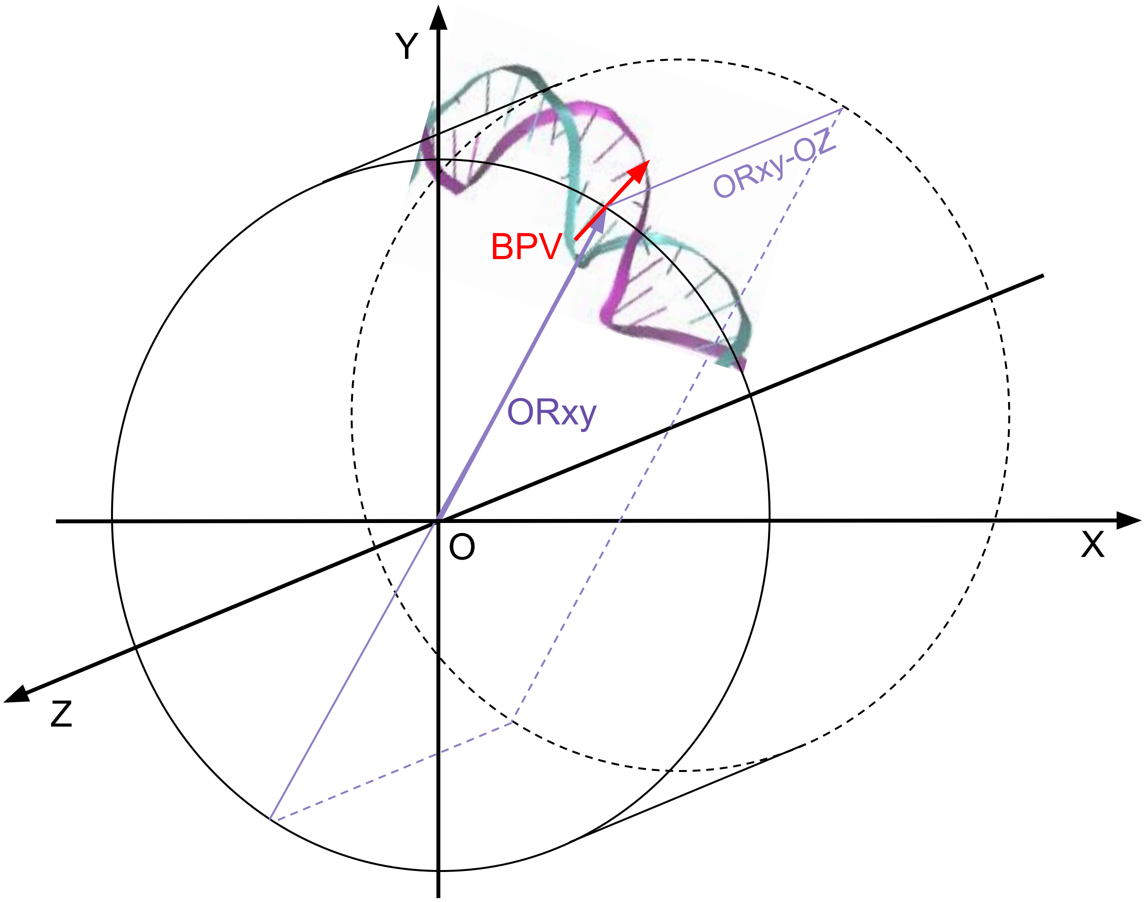


Figure SF13: Schematic description of relative twist (rTw) determination. BPV - base pair vector, ORxy - radial vector in cylindrical coordinate system, OZ - superhelical axis. rTw is defined as the angle between BPV projection onto the ORxy-OZ plane and OXY plane.

### Supplementary Methods

#### MD simulations protocols

Simulations were run using GROMACS v2018.[6] Calculations were run on the Lomonosov-2 supercomputer[7] in parallel using domain decomposition. Four to eight nodes were used, each having 14 CPU cores and one NVidia Tesla K40 GPU. Two 7-thread MPI tasks were assigned to every node. GPUs were used to calculate non-bonded interactions.[8] Simulations were performed in NPT ensemble with periodic boundary conditions (xyz). A velocity-rescaling scheme was used for temperature coupling at 300 K.[9] The extended-ensemble Parrinello-

Rahman approach was used for isotropic pressure coupling at 1 bar.[10] Verlet non-bonded cut-off scheme with grid neighbor search algorithm updated every 10 steps was used. 0.8 nm cut-off radius for van der Waals interactions with dispersion correction for energy and pressure was used. Particle Mesh Ewald (0.8 nm real space cut-off, fourth-degree PME order, 0.12 nm Fourier spacing) approach was used to account for Coulomb interactions. All bonds were constrained using the fourth-order LINCS algorithm[11] with one iteration used to correct for rotational lengthening. During MD production runs, the integration step of 2 fs was used, trajectory frames were saved every 1 ns. Production run protocol files may be found at GitHub `https://intbio.github.io/Armeev_et_al_2021/MD_production_protocol.mdp`.

To suppress potential fraying of terminal DNA base pairs during simulations, the distance between glycosidic nitrogen atoms (N1 for thymine or cytosine and N9 for adenine or guanine) was constrained using a harmonic potential with force constant of 1000 kJ/mol/nm$^2$ (approx. 4kT penalty for deviation of 1 Å). For simulations with fixed histone folds, the C$\alpha$-atoms of $\alpha$1, $\alpha$2, and $\alpha$3-helices in all histones were restrained to their original X-ray positions using a harmonic potential with force constant of 1000 kJ/mol/nm$^2$.

##### System creation and equilibration

Crystal structures were downloaded from Protein Data Bank (see PDB IDs in Table ST1). Depending on the system, we either partially truncated the flexible histone tails (see truncation sites in Figure 1e) or adjusted their conformation. In NCP$_{147}$ structure, we used UCSF Chimera[4] to adjust conformations of histone chains E, F, G, H to the conf0rmation of their pseudo-symmetry mates (chains A, B, C, D). This resulted in a more compact NCP and also allowed us to compare the evolution of tails' conformations on both sides. Crystal water was not retained in this case. Systems with truncated tails contained crystal water within 3 Å around nucleosome if available in the PDB file. Structures were positioned in the truncated octahedron boxes with the minimum distance from the border of the box to the structure of 2 nm and solvated with a TIP3P water model[12] using GROMACS command-line utilities.

$Na^+$ and $Cl^-$ ions were added to the systems to neutralize the total charge of the system and bring the ion strength to 150 mM (see details in Table ST1). Minimization was performed via the steepest descent gradient method for 10000 steps with positional restraints on heavy atoms of 500 kJ/mol/$\AA^2$. Then the system was equilibrated in five consecutive steps with gradual reduction of restraining potential: 1. 100 ps with positional restraints of 500 kJ/mol/$\AA^2$ with 0.5 fs time step; 2. 200 ps with positional restraints of 50 kJ/mol/$\AA^2$ with 2 fs time step (and further the same); 3. 200 ps with positional restraints of 5 kJ/mol/$\AA^2$; 4. 200 ps with positional restraints of 0.5 kJ/mol/$\AA^2$; 5. 200 ps of unrestrained simulations.

**Force field parameters**

Simulations were run in Amber ff14SB force field[13] with parmbsc1[14] corrections to simulate DNA. Original ions parameters were replaced with CUFIX[15] ion corrections, which contained refined Lennard-Jones parameters calibrated against experimental osmotic pressure values of DNA solutions.

**Trajectory analysis**

Custom analysis programs and pipelines were written in Python 3, integrating the functionality of GROMACS, MDAnalysis,,[16,17] VMD (visualization)[18] and 3DNA (determination of DNA base pair centers, calculation of base pair and base pair step parameters).[19] With GROMACS utilities we performed trajectory preprocessing, checked simulation parameters and the quality of simulation (energy, pressure, and some other parameters). Additional basic analysis approaches were used, such as calculations of root-mean-square deviation of atomic positions and their root-mean-square fluctuations (C$\alpha$-atoms for amino acids and P atoms for nucleotides) for amino acids sidechains and nucleotides. For DNA-histone contact analysis, atom-atom contacts were calculated as non-hydrogen atom pairs having a distance of less than 4 $\AA$.

#### Nucleosome structural elements

The main structural elements are shown in the Figure 1e. Helices are defined by taking the minimum length of alpha-helices over symmetric chains in 1KX5 X-ray structure. For alignment, C-alpha atoms of the histone folds (which are defined as $\alpha1$, $\alpha2$, and $\alpha3$-helices as in the figure) were used. Further in our analysis, we often analyzed the contacts between histone core and histone tails separately. The definition of histone tails is provided in Figure 1e, histone globular core is defined as all other parts of the histone sequence.

#### Relative twist definition

Relative twist (rTw) along the DNA sequence was determined after positioning the NCP into the nucleosome reference frame as follows (see Figure SF13). For every nucleotide pair, the base pair orientation vector (BPV) was calculated as C1'-C1' (ribose C-atoms, forming N-glycosidic bonds) vector from top to bottom DNA strand. The radial vector in the cylindrical coordinate system (ORxy) was defined as the perpendicular pointing from the nucleosome superhelical axis (OZ) to the base-pair center (BPV center). Relative twist was defined as the angle between the OXY plane (the front plane in NRF coordinates) and the projection of the BPV onto the OZ-ORxy plane. In such a definition, the parameter varies in the range from -180 to 180 degrees, but for continuity, we build the modulus of the parameter. As a result, we get a function with a period of about 10-11 base pairs and varying from 0 to 180 degrees. rTw of 90 degrees means that the base pair is parallel to the superhelical axis. Minimum corresponds to major groove orientation towards the histone core, maximum - minor groove orientation towards the core. Plotting rTw profile along the sequence allows getting information about DNA orientation, twist-defects, DNA sliding. Figure SF1 shows relative twits along the DNA in key X-ray structures (1KX5, 3LZ0, and 1AOI) and highlights the DNA sequence details and motifs.

### References


(1) Davey, C. A.; Sargent, D. F.; Luger, K.; Maeder, A. W.; Richmond, T. J. Solvent Mediated Interactions in the Structure of the Nucleosome Core Particle at 1.9 a Resolution. *Journal of Molecular Biology 319*, 1097–1113.

(2) Luger, K.; Mäder, A. W.; Richmond, R. K.; Sargent, D. F.; Richmond, T. J. Crystal Structure of the Nucleosome Core Particle at 2.8 A Resolution. *Nature 389*, 251–260.

(3) Vasudevan, D.; Chua, E. Y. D.; Davey, C. A. Crystal Structures of Nucleosome Core Particles Containing the '601' Strong Positioning Sequence. *Journal of Molecular Biology 403*, 1–10.

(4) Pettersen, E. F.; Goddard, T. D.; Huang, C. C.; Couch, G. S.; Greenblatt, D. M.; Meng, E. C.; Ferrin, T. E. UCSF Chimera–a Visualization System for Exploratory Research and Analysis. *Journal of Computational Chemistry 25*, 1605–1612.

(5) Wei, S.; Falk, S. J.; Black, B. E.; Lee, T.-H. A Novel Hybrid Single Molecule Approach Reveals Spontaneous DNA Motion in the Nucleosome. *Nucleic Acids Research 43*, e111.

(6) Páll, S.; Abraham, M. J.; Kutzner, C.; Hess, B.; Lindahl, E. In *Solving Software Challenges for Exascale*; Markidis, S., Laure, E., Eds.; Lecture Notes in Computer Science; Springer International Publishing, Vol. 8759; pp 3–27.

(7) Voevodin, V. V.; Antonov, A. S.; Nikitenko, D. A.; Shvets, P. A.; Sobolev, S. I.; Sidorov, I. Y.; Stefanov, K. S.; Voevodin, V. V.; Zhumatiy, S. A. Supercomputer Lomonosov-2: Large Scale, Deep Monitoring and Fine Analytics for the User Community. *Supercomputing Frontiers and Innovations 6*, 4–11.

(8) Abraham, M. J.; Murtola, T.; Schulz, R.; Páll, S.; Smith, J. C.; Hess, B.; Lindahl, E. GROMACS: High Performance Molecular Simulations through Multi-Level Parallelism from Laptops to Supercomputers. *SoftwareX 1-2*, 19–25.

(9) Bussi, G.; Donadio, D.; Parrinello, M. Canonical Sampling through Velocity Rescaling. *The Journal of Chemical Physics 126*, 014101.

(10) Parrinello, M.; Rahman, A. Polymorphic Transitions in Single Crystals: A New Molecular Dynamics Method. *Journal of Applied Physics 52*, 7182–7190.

(11) Hess, B. P-LINCS: A Parallel Linear Constraint Solver for Molecular Simulation. *Journal of Chemical Theory and Computation 4*, 116–122.

(12) Jorgensen, W. L.; Chandrasekhar, J.; Madura, J. D.; Impey, R. W.; Klein, M. L. Comparison of Simple Potential Functions for Simulating Liquid Water. *The Journal of Chemical Physics 79*, 926–935.

(13) Maier, J. A.; Martinez, C.; Kasavajhala, K.; Wickstrom, L.; Hauser, K. E.; Simmerling, C. ff14SB: Improving the Accuracy of Protein Side Chain and Backbone Parameters from ff99SB. *Journal of chemical theory and computation 11*, 3696–3713.

(14) Ivani, I. et al. Parmbsc1: A Refined Force Field for DNA Simulations. *Nature Methods 13*, 55–58.

(15) Yoo, J.; Aksimentiev, A. New Tricks for Old Dogs: Improving the Accuracy of Biomolecular Force Fields by Pair-Specific Corrections to Non-Bonded Interactions. *Physical Chemistry Chemical Physics 20*, 8432–8449.

(16) Michaud-Agrawal, N.; Denning, E. J.; Woolf, T. B.; Beckstein, O. MDAnalysis: A Toolkit for the Analysis of Molecular Dynamics Simulations. *Journal of Computational Chemistry 32*, 2319–2327.

(17) Gowers, R.; Linke, M.; Barnoud, J.; Reddy, T.; Melo, M.; Seyler, S.; Domański, J.; Dotson, D.; Buchoux, S.; Kenney, I.; Beckstein, O. MDAnalysis: A Python Package for the Rapid Analysis of Molecular Dynamics Simulations. pp 98–105.

(18) Humphrey, W.; Dalke, A.; Schulten, K. VMD: Visual Molecular Dynamics. *Journal of Molecular Graphics 14*, 33–38, 27–28.

(19) Lu, X.-J.; Olson, W. K. 3DNA: A Software Package for the Analysis, Rebuilding and Visualization of Three-Dimensional Nucleic Acid Structures. *Nucleic Acids Research 31*, 5108–5121.
